# A Circadian Light Regulator Controls a Core CAM Gene in the Ice Plant’s C_3_-to-CAM Transition

**DOI:** 10.1101/2025.05.03.652029

**Authors:** Noé Perron, Tam Le, Christopher Dervinis, Wendell Pereira, W. Brad Barbazuk, Matias Kirst

**Affiliations:** Plant Molecular and Cellular Biology Program, University of Florida, Gainesville, FL 32608, USA; School of Forest, Fisheries and Geomatics Sciences, University of Florida, Gainesville, FL 32603, USA; Department of Biology, University of Florida, Gainesville, FL 32608, USA; University of Florida Genetics Institute, Gainesville, FL 32608, USA

**Keywords:** Crassulacean acid metabolism (CAM), *Mesembryanthemum crystallinum*, C_3_ photosynthesis, Single-nuclei RNA sequencing, Drought tolerance, UVR8 signaling, PPCK1 regulation, HY5 transcription factor

## Abstract

Crassulacean acid metabolism (CAM) enhances drought tolerance by shifting carbon fixation to the night, improving water-use efficiency compared to C_3_ and C_4_ photosynthesis. However, the molecular regulators of CAM induction remain poorly understood. Here, we generate the first single-nucleus transcriptome atlas of a CAM species, *Mesembryanthemum crystallinum*, to resolve transcriptional dynamics at the cell-type level during the C_3_-to-CAM transition. Using snRNA-seq and a 24-hour time-course bulk RNA-seq dataset, we identify *PPCK1*, a key CAM enzyme regulator, as part of a co-expression network enriched in circadian clock genes and salt-induced pathways. We demonstrate that the ice plant HY5 (McHY5) directly activates *PPCK1*, a function absent in the C_3_ model species *Arabidopsis thaliana*. This discovery reveals a fundamental divergence in transcription factor activity between a CAM and a C_3_ species, suggesting that CAM evolution in *M. crystallinum* involved a rewiring of core regulatory elements underlying CAM. Identifying a transcription factor that directly controls a major CAM gene provides a key step toward decoding CAM regulatory architecture and opens new avenues for engineering drought-resilient crops.

## INTRODUCTION

The C_3_ photosynthetic pathway is prevalent in 90% of terrestrial plants ^1,2^, while C_4_ photosynthesis and Crassulacean acid metabolism (CAM) have evolved in about 3% and 7% of flowering plant species, respectively, as adaptations to environmental challenges like low CO_2_ or water scarcity ^2^. CAM is particularly advantageous due to its ability to fix CO_2_ primarily at night, reducing evapotranspiration and increasing water retention in leaves compared to C_3_ and C_4_ plants, which exclusively fix CO_2_ during the day ^1,3–6^. CAM plants are exceptionally drought-tolerant, achieving much higher water-use efficiency and requiring 20% of the water used by C_3_ and C_4_ plants ^6^. This enhanced water efficiency not only reduces irrigation demands and lowers associated costs, improving the economic viability of CAM crops, but also conserves critical water resources, emphasizing their important environmental and economic potential. Unlike obligate CAM species, which consistently exhibit CAM in fully-developed tissues, facultative CAM plants, such as the common ice plant (*Mesembryanthemum crystallinum*), can switch from C_3_ to CAM in response to abiotic stresses including high salinity and drought ^3,5,7–9^. This transition has been prominently studied in the ice plant, making it a model species for understanding CAM regulation ^3,7,10,11^.

Although C_4_ and CAM photosynthesis evolved similar biochemical mechanisms to concentrate CO_2_ and optimize assimilation in the Calvin-Benson-Bassham (CBB) cycle, they differ in their spatial organization. In C_4_ plants, CO_2_ concentration is compartmentalized between mesophyll and bundle-sheath cells, while in CAM plants, it occurs within a single cell type, primarily the mesophyll cells ^12^. In CAM plants, this process is regulated temporally rather than spatially, suggesting that the circadian clock controls key enzymes of the pathway. At night, CARBONIC ANHYDRASE (CA) facilitates the rapid conversion of CO_2_ to bicarbonate, which is then used by PHOSPHO*ENOL*PYRUVATE CARBOXYLASE (PPC) to fix CO_2_, resulting in the accumulation of malic acid in the vacuole ^5^. Since PPC is inhibited by malate, it is phosphorylated at night by PHOSPHO*ENOL*PYRUVATE CARBOXYLASE KINASE (PPCK), making it less sensitive to malate inhibition ^13^. *PPCK* shows a strict diel expression pattern in several CAM plants, and studies in *Kalanchoe fedtschenkoi* have demonstrated that its expression is tightly controlled, though the specific regulator remains unidentified ^3,14,15^. During the day, malate is decarboxylated in the cytosol, and the released CO_2_ is refixed by Rubisco in the CBB cycle. A major obstacle in fully understanding CAM regulation is the limited knowledge of key transcriptional regulators, such as those controlling *PPCK*, which are central to the pathway.

CAM has evolved independently multiple times across diverse plant lineages, including both monocots and dicots, on nearly every continent ^1^. This widespread and repeated evolution suggests that the transition to CAM requires only a few key genomic modifications. Specifically, it likely arises from a reconfiguration of gene expression networks, driven by changes in one or a few master regulators that alter the expression of key enzymes in response to drought ^16^. Single-cell transcriptomics has transformed molecular studies by resolving gene expression changes at the level of individual cell types, revealing regulatory shifts that traditional RNA-seq may overlook. This approach is particularly powerful for identifying master regulators of CAM, as subtle but critical transcriptional changes in key regulatory genes may be diluted in bulk RNA-seq datasets. Here, we use single-nucleus RNA sequencing (snRNA-seq) to map the transcriptional changes underlying the C_3_-to-CAM transition in *Mesembryanthemum crystallinum* at single-cell resolution. These data are complemented by a 24-hour time-course bulk RNA-seq analysis in both C_3_ and CAM-performing ice plants, providing a comprehensive view of CAM induction dynamics. In plants, the circadian clock control is not only temporally, but also spatially organized ^17^.

Unlike the centralized mammalian clock, the plant clock acts in a tissue and cell type-specific manner. By focusing on the cell-types transitioning from C_3_ to CAM, we offer new insights into the circadian clock’s association with CAM over the 24-hour cycle. ELONGATED HYPOCOTYL5 (HY5) and HY5-HOMOLOG (HYH) are conserved bZIP transcription factors that regulate growth, development, and responses to abiotic stress ^18,19^. They also bind the promoters of key circadian clock genes, including *LHY*, *CCA1*, *ELF4*, and *TOC1*, positioning them as key regulators of clock function ^18^. Their activity is regulated by CONSTITUTIVE PHOTOMORPHOGENIC 1 (COP1), which targets HY5 and HYH for degradation. However, UV-B solar radiation inhibits COP1 through interaction with UV RESPONSE LOCUS 8 (UVR8), leading to increased HY5 and HYH activity ^20,21^. Our transcriptomic analysis revealed significant co-expression of *PPCK1*, which encodes PPCK in the ice plant, with *UVR8* specifically in CAM-induced plants. Since UVR8 indirectly activates HY5 and HYH, we tested whether these transcription factors regulate *PPCK1*. Transactivation assays showed that ice plant HY5 binds to the *PPCK1* promoter from both the ice plant and *Arabidopsis thaliana*, whereas *Arabidopsis* HY5 does not bind to the *PPCK1* promoter in either species. This finding reveals a fundamental, previously unreported difference in the DNA-binding ability of a key regulator between a CAM and a C_3_ species. Our results identify HY5 as a potential regulator of *PPCK1* in CAM-performing ice plant, providing a new target for engineering drought tolerance in crops.

## RESULTS

### A high-quality ice plant genome sequence and annotation

To enable the single-nucleus transcriptome analysis of the C_3_-to-CAM transition in the common ice plant, we *de novo* sequenced, assembled and annotated the *M. crystallinum* genome. Long sequences were generated using the Oxford Nanopore Technology PromethION, resulting in reads with an N50 of 12.8kb and coverage of the genome equivalent to 175× (**Supplemental Table 1)**. Short DNA reads obtained through Illumina sequencing, achieving a coverage of 120×, were used to refine and correct errors in the draft genome assembly. The final assembly had a total length of 368 Mb distributed in 289 contigs and an assembly N50 of 7.19 Mb (n=15). A high genome completeness was confirmed by detection of a BUSCO score of 98.3%, calculated using the *embryophyta* dataset as a reference (**Supplemental Table 1**). To produce a robust annotation, the ice plant transcriptome was sequenced using PacBio Iso-Seq, generating highly accurate full-length transcripts ^22^.

To confirm the quality of the genome sequence and annotation generated in this study, a comparative analysis was conducted with the only previously published *M. crystallinum* genome (referred hereafter as IcePlant1^10^. This analysis revealed a large discrepancy in the number of annotated, protein-coding genes: 20,739 in the present study (referred to as IcePlant2) versus 24,234 in IcePlant1. Closer examination of the genes uniquely present in IcePlant1 showed that many sequences annotated as protein-coding align closely with ribosomal RNA (rRNA) sequences (**Supplemental Table 2**). In numerous instances, these rRNA sequences are replicated over a hundredfold in IcePlant1, each occurrence annotated as a unique protein-coding gene. By contrast, BLAST searches confirmed the presence of these rRNA sequences in the IcePlant2 assembly but not in the IcePlant2 coding sequences, confirming that we correctly excluded them from protein-coding annotation. In summary, the additional genes annotated in the IcePlant1 seemed to originate from the misannotation of rRNA sequences as protein-coding genes, pointing that the inclusion of full-length transcript sequencing (Iso-Seq) data supports a more accurate identification of protein-coding genes ^22,23^. The high degree of similarity between both genomes (**Supplemental Figure 1**) confirms the comprehensiveness and accuracy of the sequence generated here.

### Single-nucleus transcriptomics of the ice plant leaf deconstructs the cellular complexity of a facultative CAM organ

To determine the timing of the switch between C_3_ and CAM under our growth conditions, we conducted a daily evaluation of CAM activity in the leaves of salt-treated and well-watered plants. CAM is characterized by the nocturnal accumulation of malic acid in the vacuoles, peaking at dawn ^4^. Since total extractable leaf titratable acidity serves as a proxy for malic acid accumulation—being the predominant organic acid fluctuating over the light/dark cycle in CAM species ^5^—we measured the differences in titratable leaf acidity between salt-treated and well-watered ice plants at dawn. Our results revealed that a significant difference in acidity was observed at and after 8 days of the start of the salt treatment (t-test, p-value=5.56×10^-4^; **Figure 1A**), indicating the onset of a transition from C_3_ to CAM ^3,5^. To characterize the transcriptional profile of this transition, we isolated nuclei from 5-week-old ice plant leaves, after 8 days of salt treatment, for single-nucleus transcriptome analysis. Leaf samples were collected 30 minutes after the onset of the dark phase (Dusk Salt sample) and 30 minutes after the onset of the photoperiod (Dawn Salt sample). In addition, we isolated leaf nuclei from ice plants watered without NaCl at the same time points (Dusk Control and Dawn Control samples). Droplet-based snRNA-seq was performed for all samples using the 10× Genomics platform. Subsequent data processing yielded an integrated, normalized, batch-corrected dataset of 17,994 high-quality nuclei grouped into 17 clusters **(Figure 1B**; **Supplemental Table 12)**.

**Figure 1:**
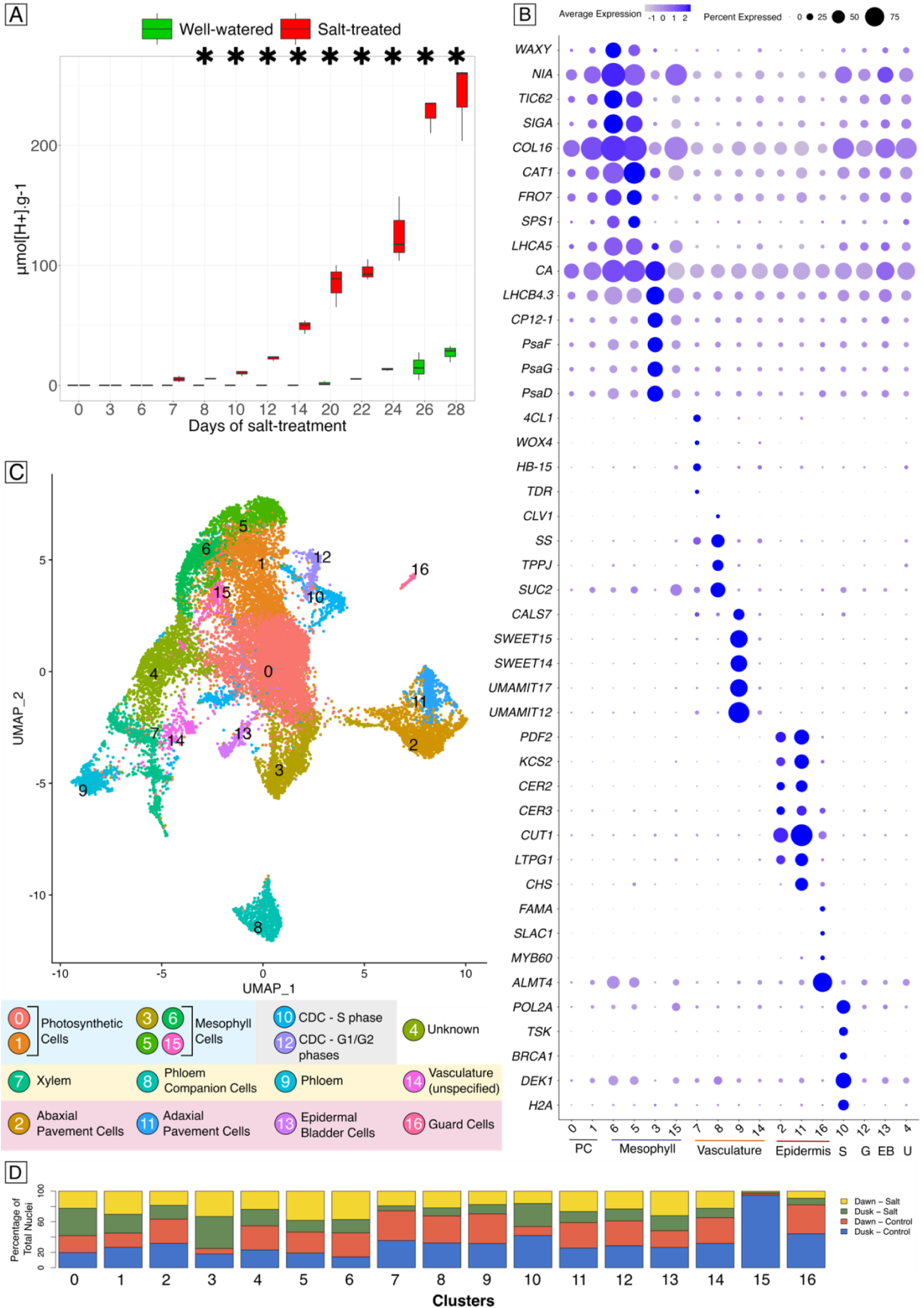
Integrated snRNA-seq data clustering and annotation. A) Boxplot representation of the evolution of titratable acidity measured in μmol[H+].g^-1^ (fresh weight) in the third leaf pair of well-watered and salt-treated ice plants. For each time-point, titration was performed on leaves from three biological replicates for each condition. Significant differences between the groups are identified by asterisks, as determined by a paired t-test (p-value < 0.05). Each boxplot illustrates the median value (black line), the first and third quartiles (the lower and upper hinges), and the extremes of the data range (the top and bottom whiskers). No outliers are depicted in this graph. B) Dotplot showing the expression of orthologs of cell type-specific markers in the integrated ice plant snRNA-seq dataset. C) UMAP clustering of the integration of all four snRNA-seq datasets (Dawn Control and Salt, Dusk Control and Salt). A total of 17,994 nuclei are divided into 17 clusters. D) Sample representation in the 17 clusters of the integrated dataset. For accurate comparison, the numbers of nuclei per cluster were adjusted based on the total number of nuclei of the sample of origin. CDC = Cell division cycle, PC = Photosynthetic Cells, S = S-phase of the cell cycle, G = G1 and G2 phases of the cell cycle, EB = Epidermal Bladder Cells, U = Unknown.

Cluster annotation was performed by examining the expression pattern of orthologs of known cell type-specific markers in the ice plant snRNA-seq dataset (see Methods; **Figure 1B&C**; **Supplemental Table 3)**. The expression patterns of vasculature-specific markers *WOX4* ^24^, *HB-15* ^25^, *TDR* ^26^, *ARF5* ^27^, *4CL1* ^28^, and *SWEET14* ^29^, *UMAMIT12* ^30^, *UMAMIT17* ^30^, *SWEET15* ^31^, and *CALS7* ^32^ were used to designate clusters 7 and 9 as xylem and phloem cells, respectively. Cluster 8 was found to comprise phloem companion cells based on markers for that cell type (*SUT1* ^33^, *FTIP1* ^33^, *TPJJ* ^34^, *SS* ^35^, *CLV1* ^36^), and cluster 14 was annotated as unspecified vasculature cells based on similarity with the expression profiles of both phloem and xylem clusters. The expression of known pavement cell (PC)-specific markers such as *CER3* ^37^, *CUT1* ^38^, *CER2* ^39^, *KCS2* ^40^, *PDF2* ^41^, and *LTPG1* ^42^ mostly localized to clusters 2 and 11. Refinement of the annotation of these clusters was facilitated by analyzing the expression of the adaxial PC-specific marker, *CHS* ^43^ which exhibited specificity to cluster 11. As a result, cluster 11 was categorized as adaxial PC and cluster 2 as abaxial PC, given the lower expression of CHS. The identification of the guard cell cluster (cluster 16) was based on the specific expression of *FAMA* ^44^, *MYB60* ^45^, and *SLAC1* ^46^. Moreover, the expression of *ALMT4*, which was shown to be limited to guard and mesophyll cells ^47^, was most pronounced in clusters 16, 5, and 6. The distinct expression of DNA replication markers such as *DEK1, H2A, BRCA1, TSK,* and *POL2A* in cluster 10 led us to classify it as nuclei within the S-phase of the cell division cycle (CDC, Suzuki et al., 2005; Joo et al., 2007; Johnson et al., 2009; Roeder et al., 2012; Pedroza-García et al., 2017).

The expression profiles of mesophyll-specific markers *SPS1* ^53^, *SIGA* ^54^, *FRO7* ^55^, *CAT1* ^56^, *TIC62* ^57^, *NIA* ^58^, *WAXY* ^59^, and *COL16* ^60^ identified clusters 5, 6, and 15 as encompassing mesophyll cells. The expression of photosynthesis-associated genes previously found to be upregulated in the mesophyll such as *LHCA5* ^61^, *CA* ^62^*, PsaD* ^63^*, PsaG* ^64^*, PsaF* ^65^*, CP12-1* ^66^, and *LHCB4.3* ^67^ facilitated the identification of clusters 3 as comprising mesophyll cells as well ^42,68^. Clusters 0 and 1 exhibited a lower level of gene expression associated with mesophyll-specific markers compared to clusters 3, 5, 6, and 15. However, both clusters were primarily characterized by the expression of genes involved in photosynthesis, leading us to annotate these clusters as containing a mix of photosynthetic cells. This pattern aligns with observations from other snRNA-seq analyses of leaves, where the largest clusters are often characterized by a high abundance of photosynthesis-related genes but not a definitive cell type ^33,69–72^.

Upon annotating most clusters using orthologs of established cell type-specific markers, the identities of clusters 4, 12 and 13 remained ambiguous. For cluster 12, the prevalence of genes associated with auxin metabolism, growth, and cell elongation hinted at cells characterized by robust growth, potentially resembling G1/G2-phase cells in the cell division cycle (CDC). For example, genes known to be linked to cell elongation and growth, such as *LNG2* ^73^, *ARF19* ^74^, and *IAA7* ^75^ were found to be specific to cluster 12 (average log2 fold change compared to their expression in other clusters > 2 ; **Supplemental Table 4**). Thus, cluster 12 was defined as a cell elongation/growth cluster. The same strategy was employed to gain deeper insights into the unique characteristics of cluster 13. This analysis unveiled a predominance of transcripts related to ion exchange, homeostasis, transport, and storage functions. Notably, these features aligned with the transcriptional signature of epidermal bladder cells. These specialized trichomes play a crucial role in sequestering excessive salts in *M. crystallinum*, enabling the plant to maintain normal function under high salinity conditions, a pivotal process for salt tolerance ^76^. Consequently, we designated cluster 13 as a putative epidermal bladder cell cluster. Finally, cluster 4 lacked a clear transcriptional identity – after investigation of the genes specifically expressed in this group, it was annotated as “unknown”.

### Mesophyll cells exhibit high transcriptomic plasticity during the C_3_-to-CAM switch

The categorization of mesophyll cells into four distinct clusters (3, 5, 6 and 15) prompted us to investigate the transcriptional differences underpinning such heterogeneity. An examination of each cluster’s sample composition (proportional to the total number of nuclei per sample) showed that 41.9% of nuclei in cluster 3 were derived from the Dusk – Salt sample. Similarly, the vast majority of nuclei in cluster 15 originated from the Dusk – Control sample. On the other hand, a predominant fraction of nuclei in clusters 5 and 6 were from the Dawn – Salt sample, at 38.1% and 36.9%, respectively (**Figure 1D)**.

Using a non-parametric Wilcoxon rank sum test, we then compared transcript levels between nuclei from salt-stressed samples (Dawn and Dusk Salt) to those from well-watered samples (Dawn and Dusk Control) within each mesophyll cluster (specifically, clusters 3, 5, 6, and 15; **Supplemental Table 5**). Genes identified as differentially expressed (DE, with an adjusted p-value < 0.05) were categorized according to their average log2 fold-change as follows: “highly upregulated” (log2 fold-change > 1.5), “upregulated” (1.5 > log2 fold-change > 1), and “moderately upregulated” (1 > log2 fold-change > 0). DE genes with a log2 fold-change below 0 were classified as “downregulated.” Based on this classification, we observed a high upregulation of transcripts for key CAM pathway genes—such as *PPC1*, *NADP-ME*, and *PPDK*—which are known to peak in expression at dusk ^3,77^. This upregulation was specifically detected in cluster 3 nuclei from salt-treated samples, a trend not observed in other mesophyll clusters (**Supplemental Table 5**). These findings indicate that cluster 3 contains CAM mesophyll cells in their dusk transcriptional state.

Three copies of *PPCK1* were found in our sequenced genome of *M. crystallinum*. The expression patterns and cell-type specificity of two of these copies, *PPCK1.1* and *PPCK1.2*, were highly similar, while expression of the third copy was not detected in our dataset. *PPCK1.1* and *PPCK1.2* occur in tandem in *M. crystallinum* genome, suggesting a recent gene duplication, and are known to peak in expression just before dawn and show minimal expression at dusk in the ice plant ^3,77^. While these genes were not upregulated in cluster 3, they were highly upregulated in the nuclei of clusters 5 and 6 from salt-treated samples **(Figure 2A,B,C, Supplemental Table 5**). Such patterns, as well as the observations made previously, indicated that cluster 3 was indeed enriched with dusk CAM mesophyll cells, while clusters 5 and 6 predominantly harbored dawn CAM mesophyll cells. The salt-induced stress did not lead to any variation in CAM enzyme encoding transcripts within cluster 15 **(Figure 2A,B,C)**. This, together with the observation that genes DE in cluster 15 are known to be exclusively expressed at night (e.g., *LEA5* ^78^, *TOC1* ^79^, **Supplemental Table 7**) enabled its annotation as “dusk C_3_ mesophyll cells”. To further support our temporal annotation of mesophyll clusters, we analyzed the expression patterns of pivotal regulators of the plant circadian clock (**Figure 2D**). Genes crucial for sensing dawn and photoperiod, such as *LHY*, *UVR8*, *PRR9*, *LNK1, LNK2*, *RVE1*, *HY5,* and *ZTL* ^18,19,80^ showed pronounced expression in clusters 5 and 6. In contrast, the evening-specific gene *TOC1* and other components of the evening complex of the core circadian clock network, including *LUX* and *ELF4* ^79^, exhibited elevated expression specifically in the C_3_ cells of cluster 15. These genes did not present a similarly heightened expression in the CAM cells of cluster 3. This aligns with the documented shift in core circadian clock evening complex gene expression seen in *Kalanchoë fedtschenkoi*, a constitutive CAM species ^81^.

**Figure 2:**
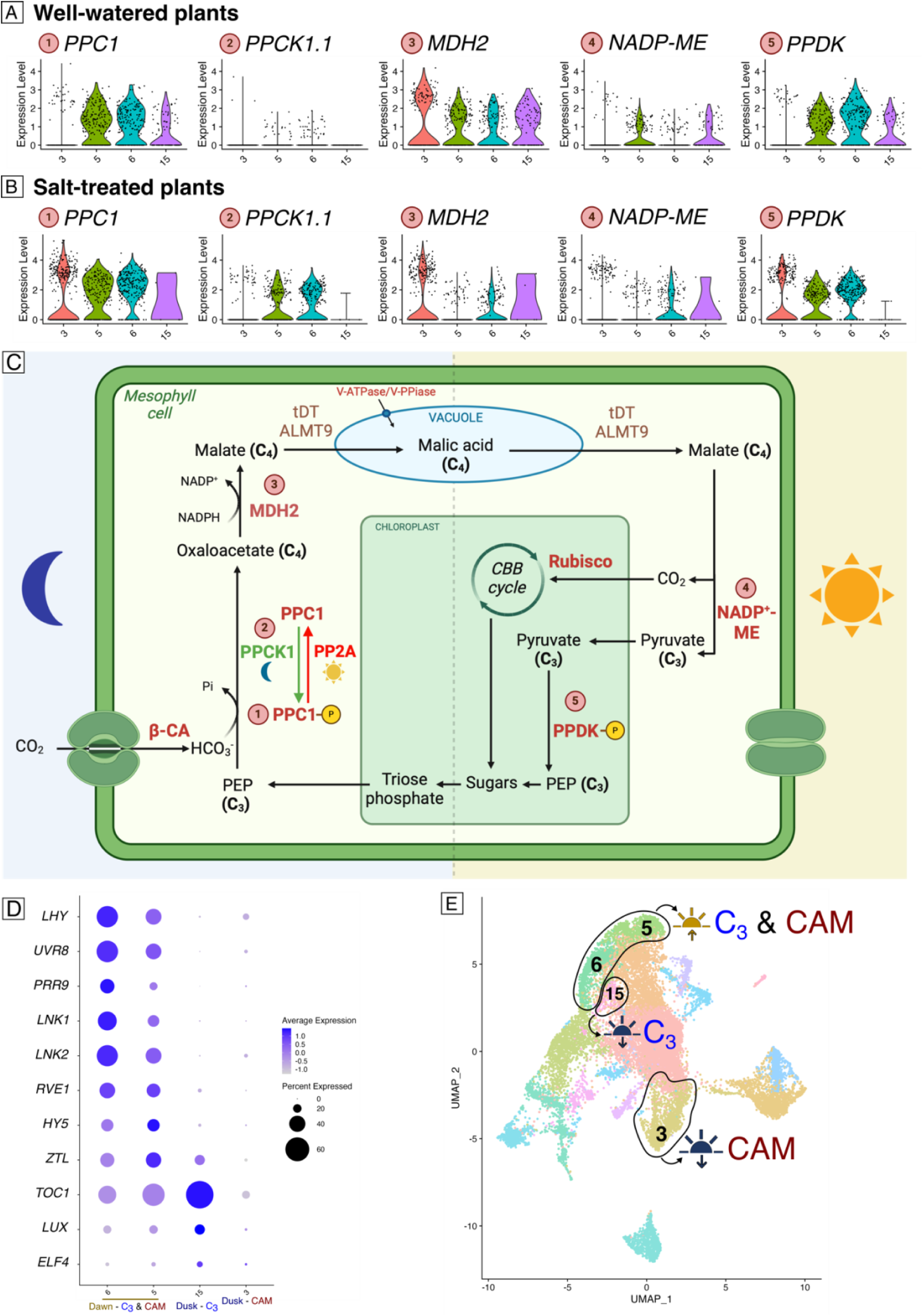
Expression profiles of CAM and circadian clock genes in the snRNA-seq data. A) Violin plots depicting cluster-specific expression profiles of CAM genes in the mesophyll cells of the well-watered samples (Dawn Control and Dusk Control). B) Violin plots depicting cluster-specific expression profiles of CAM genes in the mesophyll cells of the salt-treated samples (Dawn Salt and Dusk Salt). The two copies of PPCK1 expressed in our dataset display nearly identical expression profiles. Therefore, only the first copy of *PPCK1* in the ice plant genome (*PPCK1.1*) was represented in this graph. The numbers in front of the gene names in Figure 2A&B correspond to the numbers in Figure 2C. C) Overview of the main biochemical events of CAM during the 24-hour diel cycle in CAM-performing ice plant mesophyll cells. During the night, atmospheric CO_2_ is converted to HCO ^-^ by BETA-CARBONIC ANHYDRASE (β-CA). This product is then added to phospho*eno*lpyruvate (PEP) by PEP CARBOXYLASE 1 (PPC1) to generate oxaloacetate (OAA). PPC1 is activated at night via phosphorylation by PPC KINASE 1 (PPCK1), and deactivated during the day via dephosphorylation by PROTEIN PHOSPHATASE TYPE 2A (PP2A, Carter et al., 1990). OAA is rapidly reduced in the cytosol by MALATE DEHYDROGENASE 2 (MDH2). Malate is then transported into large vacuoles where it is stored as malic acid for the remainder of the dark phase. During the daytime, stomata close to minimize water loss by transpiration, and malate is transported out of the vacuole. Subsequently, it is decarboxylated in the cytosol by NAD(P)-MALIC ENZYME (NADP-ME), or NAD-MALIC ENZYME, providing CO_2_ for daytime fixation via Rubisco in the Calvin-Benson-Bassham (CBB) cycle. The pyruvate product of decarboxylation is converted to PEP by PYRUVATE, PHOSPHATE DIKINASE (PPDK) in the chloroplast ^83^. D) Cluster-specific expression of a selection of regulators of the plant circadian clock in the cells annotated as mesophyll. E) Schematic representation of the annotation of each mesophyll cell cluster in the UMAP space.

Our clustering analysis effectively distinguished between C_3_ and CAM mesophyll cells at dusk. However, no such separation was achieved at dawn, with both cell states blending into clusters 5 and 6. This suggests that the transcriptional differences between CAM- and C_3_-performing cells at dawn are less pronounced during the early stages of CAM induction, with most genes being similarly expressed in both cell states. Therefore, more refined computational analyses are needed to identify the key differences between the two pathways at dawn, as we demonstrate next.

### *PPCK1* expression is correlated to the circadian clock in CAM-induced samples

After identifying clusters enriched with nuclei from dawn or dusk samples (**Figures 2D & 2E**), we performed gene co-expression network inference analyses at key points in the CAM cycle. We applied high-dimensional weighted gene co-expression network analysis (hdWGCNA, Morabito et al., 2023) to the integrated dataset, resulting in 10 gene modules with distinct cluster affinities (**Figure 3A**). Genes within each module were ranked according to their eigengene-based connectivity (kME), which quantifies their association with a specific module. A kME value of 0 indicates no connection, values approaching 1 indicate a strong positive association, and values approaching -1 indicate a negative association (**Supplemental Table 6**, **Figure 3A, Supplemental Figure 2**). To identify potential CAM regulators, we focused our downstream analyses on modules containing known CAM enzymes. This approach identified Modules 1, 2, 7, and 9 as containing the main CAM enzymes with varying levels of connectivity (**Figure 3A**). Since some of these modules might include networks unrelated to CAM induction, we further analyzed the proportion of genes moderately to highly upregulated by salt treatment in the snRNA-seq dataset (log2 fold-change > 0.5, adjusted p-value < 0.05, **Supplemental Table 7**) within each module. This analysis highlighted Module 2 as the most populated by genes whose expression increased in conjunction with the CAM induction process, with 106 out of the 168 salt-responsive upregulated genes included in this module (**Figure 3B**). The presence of *PPCK1.1* and *PPCK1.2* in Module 2 (kME = 0.43 and 0.40, respectively) and the high connectivity of circadian clock genes at dawn within this module (e.g., *LHY* (kME = 0.86), *UVR8* (kME = 0.79), *LNK2* (kME = 0.69), *RVE8* (kME = 0.61); **Supplemental Table 6**, ^85^ indicated a potential association between CAM and circadian clock regulation in the morning in *M. crystallinum*.

**Figure 3:**
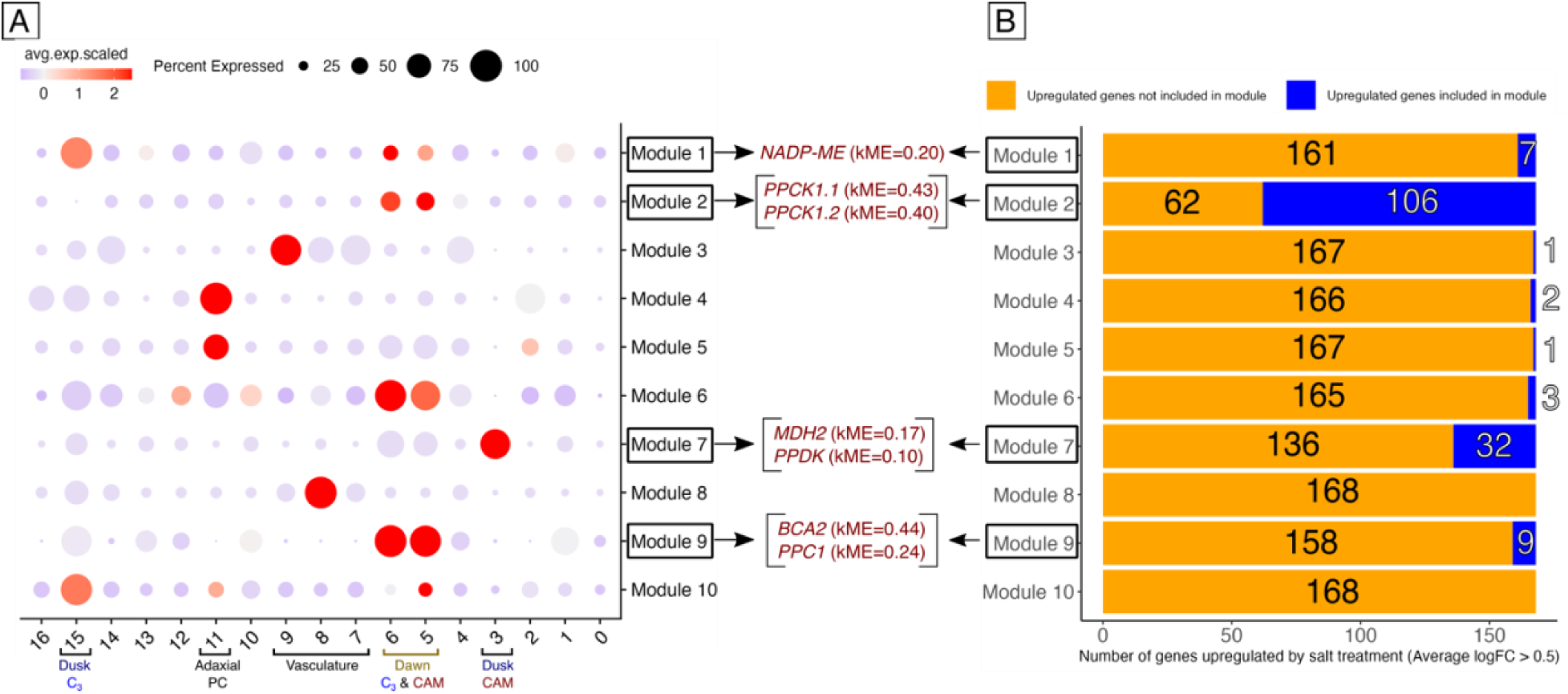
Overview of the results from the hdWGCNA analysis. A) Dot plot illustrating cluster-specific expression of genes contained in hdWGCNA modules from snRNA-seq data. Modules 1, 2, 7, and 9 include key enzymes of the CAM cycle. B) Bar plot illustrating the distribution of 168 genes upregulated by salt treatment in our snRNA-seq data (average logFC > 0.5) across the different hdWGCNA modules. The orange bars represent the number of upregulated genes not included in each module, while the blue bars indicate the number of upregulated genes included within the module.

### Bulk RNA-seq analysis strengthens the prediction of a *PPCK1* co-expression network in CAM-performing *M. crystallinum*

To obtain further supporting evidence for a regulatory network including *PPCK1* and clock genes, we sought evidence of similar co-expression links in an independent experiment with fully CAM-induced ice plants. Twenty-one days of salt treatment resulted in a significantly advanced stage of CAM induction (**Figure 1A**). Thus, we conducted a time-course bulk RNA-seq experiment with six time points over a 24-hour cycle in ice plants after 21 days of salt treatment. The same experimental design was applied to well-watered, C_3_-performing plants of the same age as controls (**Figure 4A**). We utilized the newly generated datasets to investigate active co-expression networks at specific times using the maSigPro package, designed to identify genes with changing expression profiles across conditions (here, time and treatment) in time-course transcriptomic experiments ^86^. The algorithm identified four clusters, each with characteristic expression patterns (**Figure 4B**). Notably, cluster 1 predominantly consisted of genes with clear diel rhythms, peaking between the end of the dark phase and dawn, with low expression levels during the light phase and at dusk (**Figure 4B**). Additionally, cluster 1 was enriched with genes upregulated by salt treatment. Exploration of this network revealed that both *PPCK1.1* and *PPCK1.2* had high connectivity (R^2^ = 0.84 and R^2^ = 0.78, respectively; **Supplemental Table 8**), validating our previous findings that *PPCK1* is part of a large co-expression network at dawn in CAM-performing ice plants. To identify genes most consistently associated with *PPCK1* in CAM-induced ice plants, we searched for those that: 1) are in the same module as both *PPCK1* genes in our snRNA-seq data, 2) are in the same cluster as both *PPCK1* genes in our bulk RNA-seq data, and 3) are upregulated in salt-treated samples after 8 and 21 days of treatment. This search resulted in five genes meeting all criteria: *ABI FIVE BINDING PROTEIN 2* (*AFP2*), *CBL-INTERACTING SERINE/THREONINE-PROTEIN KINASE 11* (*CIPK11*), *IQ-DOMAIN 23* (*IQD23*), *ULTRAVIOLET-B RECEPTOR 8* (*UVR8*), and *SNRK2-INTERACTING CALCIUM SENSOR* (*SCS*, **Supplemental Table 9**). Examination of the specific expression profiles of these five genes, along with the *PPCK1* genes, revealed highly similar transcriptional patterns (**Figure 4C**). All seven genes (*AFP2*, *CIPK11*, *IQD23*, *UVR8*, *SCS*, *PPCK1.1* and *PPCK1.2*) demonstrated induction upon salt stress and followed the same rhythmic transcription pattern in CAM-performing ice plants. Notably, the *A. thaliana* orthologs of several of these genes are involved in signaling pathways relevant to CAM induction. Both *IQD23* and *SCS* are involved in calcium signaling, a process known to promote the transcription of essential CAM genes in the ice plant, such as *PPC1* ^87–90^. *CIPK11*, also activated by calcium signaling, regulates stomatal aperture in response to multiple abiotic stresses ^91–93^. On the other hand, *AFP2*, a negative regulator of ABA signaling, plays a role in the salt stress response in *A. thaliana*, excluding a role specific to CAM-induction ^94^. *UVR8*, a UV-B receptor and key circadian clock regulator, was the only clock-associated gene consistently co-expressed with PPCK1 ^95,96^. Given the established role of circadian regulation in ensuring proper cycling of CAM enzymes ^97^, our data suggest that *UVR8* may contribute to CAM regulation in the ice plant through *PPCK1*. Further supporting this hypothesis, *UVR8* was not upregulated by salt stress in *A. thaliana* (log fold-change > 0.5, ^98^) while it was induced in CAM-performing ice plants. Interestingly, among all genes co-expressed with *PPCK1*, *UVR8* showed the largest shift in its daily expression pattern between well-watered and salt-treated conditions—its lowest expression occurred at ZT8 in well-watered plants but was delayed to ZT16 in salt-treated, CAM-performing plants (**Figure 4C**).

**Figure 4:**
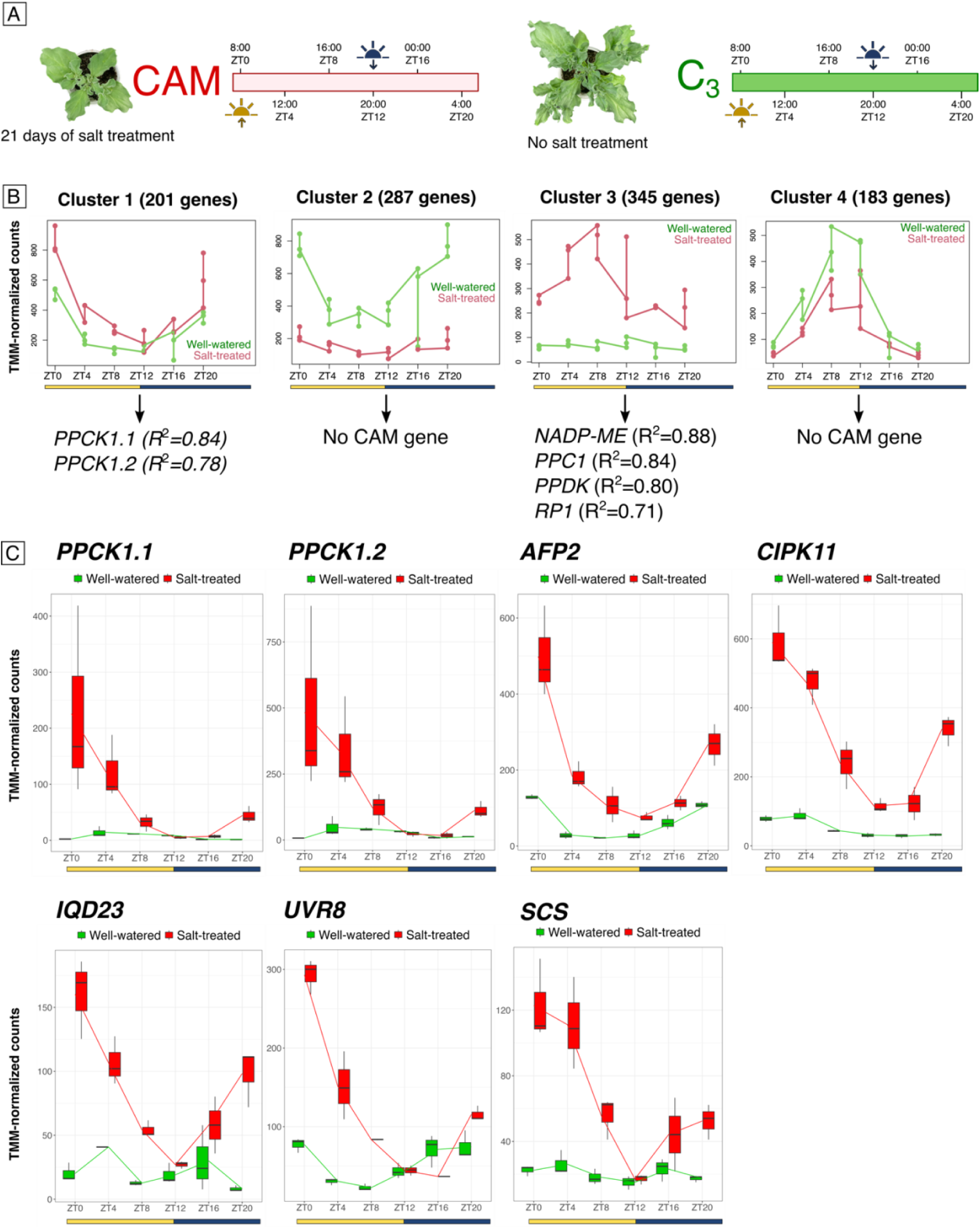
Bulk RNA-seq experimental design and overview of co-expression network analysis results. A) Schematic representation of the time-course bulk RNA-seq experimental design. Seven-week-old ice plant leaf samples were collected from both CAM-performing (21 days of salt treatment) and C_3_-performing (no salt treatment) ice plants every 4 hours over a 24-hour cycle. Tissue collection commenced at 8:00 AM (dawn, ZT0) and concluded at 4:00 AM the following day (ZT20), 4 hours before dawn. The timeline indicates the sampling points and corresponding Zeitgeber times (ZT), with light and dark phases denoted by sun going up and sun going down symbols, respectively. B) Time-series gene expression networks in response to salt treatment and well-watered conditions. The four graphs illustrates gene expression profiles over the 24-hour period as described in A) for four gene networks identified using the maSigPro R package. Each network’s expression pattern is shown for plants under well-watered (red) and salt-treated (green) conditions at six time points. The yellow bar represents the light period, while the dark bar represents the dark period. The CAM genes identified in each network, along with their correlation values, are displayed at the bottom of the graphs. C) Expression profiles of *PPCK1.1, PPCK1.2*, *AFP2*, *CIPK11, IQD23, UVR8,* and *SCS* over the 24-hour cycle described in A) in C_3_-performing (green boxes) and CAM-performing (red boxes) ice plants. The data are represented as TMM-normalized counts with error bars indicating the standard error of the mean. All three genes exhibit peak expression at dawn (ZT0) and a gradual decline throughout the day in salt-treated (CAM) plants, while their expression remains low in well-watered (C_3_) plants.

### McHY5 controls *PPCK1* expression in *M. crystallinum*

UVR8 influences the circadian clock by stabilizing HY5, a transcription factor (TF) which regulates core clock components by binding ACGT elements, including A-box (TACGTA), C-box (GACGTC), G-box (CACGTG), and Z-box (ATACGGT) motifs in target gene promoters ^99,100^. To explore the potential link between UVR8, PPCK1, and the circadian clock, we analyzed PPCK1 promoter regions (defined as 2 kb upstream of the start codon) in the ice plant genome. This analysis identified an A-box motif at -937 bp in the *PPCK1.1* promoter and two A-box motifs at - 331 and -671 bp in the *PPCK1.2* promoter, motifs that are absent from the *Arabidopsis thaliana* homolog *AtPPCK1* (*AT1G08650*). To assess whether HY5 induces *PPCK1* expression in the ice plant, we performed transactivation assays in *Nicotiana benthamiana* leaves using *PPCK1.1* and *PPCK1.2* promoters, as well as the *AtPPCK1* promoter to compare a CAM and a C_3_ species. LUCIFERASE activity was measured in the presence and absence of ice plant McHY5 (Mcr-005278), McHYH (Mcr-009873), and their *Arabidopsis* orthologs, AtHY5 and AtHYH. Of these, only McHY5 significantly activated LUCIFERASE expression compared to the control (**Figure 5A, Supplemental Table 13**). Notably, McHY5 induced LUCIFERASE activity regardless of the *PPCK1* promoter used, whereas AtHY5 failed to activate expression, even when paired with the *AtPPCK1* promoter. This finding highlights a fundamental difference in HY5’s regulatory function between ice plant and *Arabidopsis*, suggesting that McHY5 drives *PPCK1.1* and *PPCK1.2* induction in CAM-performing ice plant. Interestingly, none of the promoters exhibited significant LUCIFERASE activity when both McHY5 and McHYH were present (**Figure 5A**). The ability of McHY5 to significantly activate LUCIFERASE expression driven by the *AtPPCK1* promoter, despite the absence of the identified ACGT-containing elements in its sequence, suggested that McHY5 may have altered binding specificity compared with AtHY5. Furthermore, the observation that *McHY5* transcript levels are significantly lower in salt-treated ice plant leaves compared to well-watered controls (log fold-change < -1, p-value < 0.05; **Supplemental Figure 3**) supports the idea that *UVR8* upregulation, rather than increased *McHY5* transcription, is the primary driver of *PPCK1* induction in CAM-performing plants. This also implies that McHY5 abundance is likely regulated post-transcriptionally—at the protein level—through enhanced UVR8-mediated inhibition of COP1 activity.

**Figure 5:**
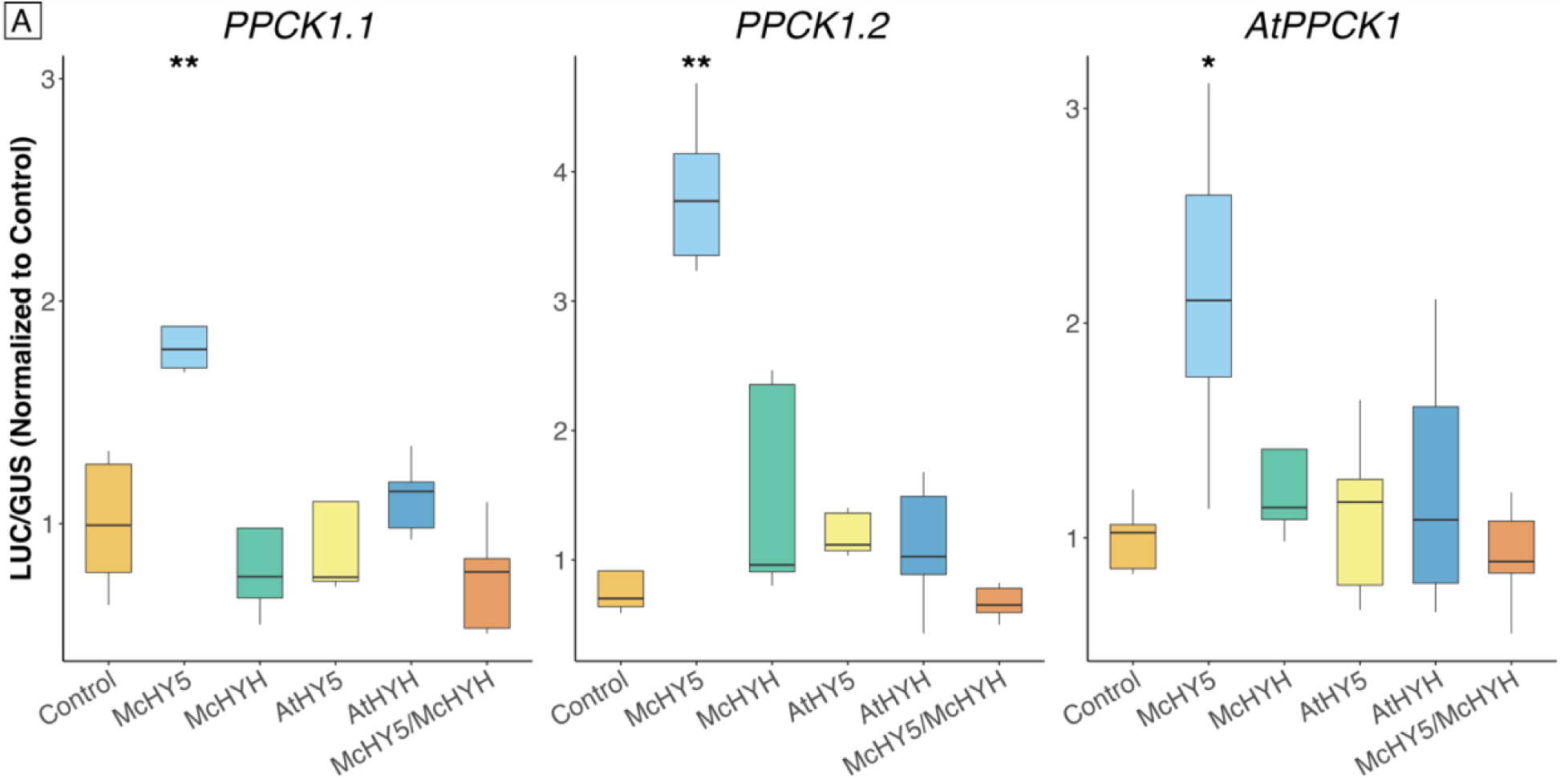
McHY5 controls *PPCK1* expression in *M. crystallinum*. Transactivation assays in *Nicotiana benthamiana* leaves reveal that McHY5 significantly induces *PPCK1.1*, *PPCK1.2*, and AtPPCK1 promoter activity, as measured by LUC/GUS luminescence normalized to the control. McHY5 activation of ice plant *PPCK1* promoters was the most pronounced, with a significant increase relative to the control (p < 0.01, Wilcoxon test). McHY5 also significantly upregulated AtPPCK1 (p < 0.05, Wilcoxon test), while McHYH, AtHY5, and AtHYH failed to induce *PPCK1* expression above control levels. Co-expression of McHY5 and McHYH did not enhance promoter activity. Data are presented as boxplots displaying the median, quartiles, and range; statistical significance is indicated (p < 0.05 (*); p < 0.01 (**)). Five biological replicates were performed for each TF/target promoter combination.

## DISCUSSION

The first single-nucleus transcriptome atlas of a CAM species provides a foundation for elucidating the underlying molecular mechanisms behind cellular function and diversity in an organism capable of thriving under extreme salinity and drought conditions. Despite the inherent recalcitrance of the ice plant to genetic transformation ^101^, hindering the direct validation of cell cluster identity, the employment of orthologs to well-established cell type-specific markers enabled us to generate a robust cluster annotation (**Supplemental Table 3**). The consistent and highly specific expression patterns of these markers within our identified cell clusters further reinforced the reliability of their annotations (**Figure 1B**). Our experimental design facilitated the computational isolation of guard cells under distinct environmental conditions. However, the limited number of guard cell nuclei in our dataset (145 across all samples) prevented the comprehensive exploration of transcriptional changes specific to them during the C_3_-to-CAM transition.

A major feature of the snRNA-seq data analysis is the identification of clusters enriched in C_3_ or CAM cells, which enables high resolution comparisons among these forms of photosynthesis. Additionally, identifying clusters containing mesophyll cells in their dusk and dawn transcriptional states facilitated further gene expression studies at the key transition phases of CAM (**Figure 2A-E**). High-dimensional co-expression network analyses using hdWGCNA revealed that *PPCK1.1* and *PPCK1.2*, which encode the kinase responsible for the phosphorylation of PPC1, the nighttime carboxylase of CAM ^102^, was strongly associated with key regulators of the core circadian clock in the morning, within a network largely characterized by the presence of salt-induced genes (**Figure 2C**, **Figure 3A&B**). Subsequent time-series gene expression studies during advanced CAM induction identified five candidate regulatory genes co-expressed with *PPCK1.1* and *PPCK1.2*, among which *UVR8* plays a critical role in the UV-B signaling pathway in plants and is a known mediator of the plant circadian clock ^21,96^. Given that *UVR8* shares a similar diel expression pattern with *PPCK1* in CAM-induced ice plants, and that it indirectly activates *HY5* and *HYH*, we tested whether these TFs could induce *PPCK1* expression in the ice plant. Our findings identify McHY5 as a regulator of *PPCK1* expression in *M. crystallinum*, suggesting its functional divergence from *Arabidopsis* HY5. Unlike AtHY5, McHY5 strongly activated *PPCK1* expression, even in the absence of conserved ACGT-containing elements in the *AtPPCK1* promoter, indicating an altered regulatory function. This discovery reveals a fundamental divergence in transcription factor activity between a CAM and a C_3_ species, suggesting that CAM evolution in *M. crystallinum* involved regulatory network modifications rather than novel gene invention. Instead of requiring the emergence of new genes, the transition to CAM may have been facilitated by changes in gene regulation, allowing pre-existing transcription factors to integrate CAM genes into their regulatory networks.

The substantial upregulation of *PPCK1* under salt stress in CAM-induced ice plants aligns with increased *UVR8* expression and its downstream effects on COP1, which stabilizes HY5. The observed peak in *COP1* expression just before dusk—when *UVR8* and *PPCK1* levels are lowest (**Supplemental Figure 3**)—suggests a tightly regulated temporal mechanism linking light signaling to CAM metabolism. This integration may have evolved through modifications in McHY5, allowing it to regulate a key enzyme of the CAM dark phase in response to environmental cues. Whether additional transcription factors contribute to *PPCK1* regulation remains an open question. Despite their functional differences, sequence alignment revealed that McHY5 and AtHY5 share a highly conserved DNA-binding domain, with only three residue differences, including a semi-conservative asparagine-to-serine substitution at position 9 (**Supplemental Figure 4**). This suggests that McHY5’s altered function may arise from changes in protein interactions rather than DNA-binding affinity, or from the presence of an atypical HY5-binding motif in the *PPCK1* promoters of the ice plant.

Together, these findings suggest that McHY5 has been co-opted to regulate *PPCK1* in CAM-performing ice plants, likely through changes in transcription factor function, promoter accessibility, or upregulation of *UVR8* under drought-like conditions. This regulatory shift may represent a key evolutionary step in the integration of CAM into the plant circadian network.

## MATERIALS AND METHODS

### Genome sequencing

High molecular weight genomic DNA was extracted using 40g of fresh tissue following a protocol consisting of nuclei isolation ^103^ followed by SDS based isolation ^104^ and NaCl cleanup ^105^. Long-reads were sequenced using the Oxford Nanopore Technology (ONT, UK) PromethION platform, and the library was constructed according to the respective manufacturer’s protocols (ONT). The Covaris g-TUBE was used to shear genomic DNA into fragments between 6 and 20 kb. Fragments from the nanopore library were size selected at 12 kb using a Electrophoretic Lateral Fractionator (ELF). Short reads were sequenced with the Illumina (USA) NovaSeq 6000 platform, and the library was constructed according to the respective manufacturer’s protocols (NEBNext® Ultra™, New England Biolabs, USA). The ELF assay, and PromethION and NovaSeq6000 sequencing were carried out at the NextGen Sequencing Core, Interdisciplinary Center for Biotechnology Research (ICBR, RRID:SCR_019152) at the University of Florida.

### Genome assembly and polishing

The output reads from the PromethION were assembled using FLYE ^106^, an assembler designed for nanopore reads, using a reduced coverage of 50X by selecting the longest fragments. The resulting assembly was polished using Pilon to correct local misassemblies and errors ^107^. A Benchmarking Universal Single-Copy Orthologs (BUSCO, ^108^) score using the *embryophyta* dataset was calculated to evaluate genome completeness.

### Transcriptome sequencing

The transcriptomic profile of the ice plant was generated utilizing PacBio Iso-seq ^22^. RNA was extracted from 57 plants at diverse developmental stages, ranging from seedling to flowering, grown under two distinct conditions (well-watered and subjected to salt stress), during both diurnal and nocturnal periods. This extraction included leaves, stems, and roots, and followed a protocol previously established ^109^. The subsequent IsoSeq analysis was conducted on the PacBio SEQUEL IIe platform at the ICBR NextGen Sequencing core. The cDNA was size selected and separated into two groups: ≤ 2.5 kb and > 2.5 kb in length. Fragments larger than 2.5 kb were subjected to reamplification, and subsequently, cDNA from both size categories were pooled equimolarly in preparation for sequencing. A single SMRT (Single-molecule real-time sequencing) cell run was performed with a movie time of 30 hours. PacBio sequences were processed through the IsoSeq3 workflow available in the PacBio SMRT tools v.11 package.

### Genome annotation

*de novo* genome annotation was carried out using MAKER2 ^110^, and gene discovery was supported by the PacBio Iso-Seq data. Ice plant repeat sequences were identified within the draft genome assembly using Repeatmodeler v2.0 ^111^. 2,725 repeat sequences ranging in size between 30bp and 14,057bp were identified by repeatmodeler and subsequently used by MAKER2 to mask regions of repetitive DNA. Three rounds of MAKER2 were completed to acquire the ice plant protein-coding gene annotation. The final high-quality annotation set included all gene models derived from the third round of MAKER2 that had an Annotation Edit Distance (AED) of less than 1.0, indicating overlap between *ab initio* predictions and empirical evidence. This last annotation also included all purely *ab initio* exhibiting evidence of one or more PFAM domains.

### Genome comparison

Genome-wide alignments and dot plots were generated with the D-GENIES web application ^112^ using the Minimap2 ^113^ v2.24 software package. Analysis of the genes present in the contigs unique to IcePlant1 was performed using BLASTN alignment against the nucleotide NCBI database with an e-value threshold of 1x10^-05^.

### Plant material growth conditions for snRNA-seq

Seeds of *M. crystallinum* were germinated in moistened soil, and seedlings were transferred to 946mL containers one week after germination. Plants were placed in a growth chamber under 150 μmol m^−2^ s^−1^ white lights to avoid CAM induction due to high photosynthetic photon flux density. The chamber had a 12h/12h light/dark cycle with temperatures of 26°C during the light phase and 18°C during the dark phase, and relative humidity of 50%. After transplantation, plants were watered daily with 50mL of 0.5X Hoagland’s solution. Four weeks after sowing, a group of plants was watered daily with 50mL 0.5X Hoagland’s solution containing 0.5M NaCl to induce CAM.

### Titratable acidity

Leaf nocturnal acidification was measured in both well-watered and salt-treated plants following a previously described method ^3^. Leaves from the third leaf-pair were collected at the start of the light period (hour 0), and 2g of fresh mass per sample were homogenized in 80% methanol. The leaf tissue homogenate was then titrated against 5 mM NaOH until reaching pH 7 using a pH meter. For each condition, leaves were collected from three biological replicates at each time-point for a total of six biological samples per time-point.

### Sample collection for snRNA-seq

For snRNA-seq analysis, the third pair of leaves was collected from 36-day old plants that were either well-watered or salt-treated for 8 days. Samples were collected during both the light (after 30 minutes into the 12h light period), and dark phases (after 30 minutes into the 12h dark period), resulting in four samples at 36 days. The same collection procedure was applied to the unique 28-day old sample (Day 0 of salt-treatment). Tissue collection was performed immediately before entering a walk-in cold chamber at 4°C, followed by nuclei isolation.

### Nuclei isolation from *M. crystallinum* leaves for snRNA-seq

Nuclei isolation was performed following a previously described method ^114^. All buffers and materials used were pre-cooled at 4°C, and the entire procedure was conducted at the same temperature. For the dark-phase samples, the procedure was performed under darkness, with only an overhead green light used as the light source. Entire leaves were fragmented on a glass plate in 200 μL of Nuclei Isolation Buffer (NIB, ^114^) containing 0.5 U/mL Protector RNase Inhibitor (Sigma Aldrich) using a razor blade. Nuclei in solution were then transferred to a 50 mL tube containing NIB and placed onto a rotating shaker for 5 minutes. Homogenates were then successively filtered through miracloth (Calbiochem), and then 40 μm and 20 μm filters. Samples were centrifuged at 600g for 5 minutes and washed two times using NIB WASH buffer ^114^. The resulting pellets were resuspended in 450 μL NIB WASH, and filtered one last time using 40 μm filters.

### Nuclei sorting

Following nuclei isolation, the samples were stained with 5 μg/mL DAPI and incubated for 5 minutes at room temperature. The nuclei were then isolated from the cell debris in suspension using Fluorescence Activated Nuclei Sorting (FANS) utilizing the BD FACSAria™ IIU/III at the ICBR Flow Cytometry Core at the University of Florida (RRID:SCR_019119). The nuclei sorting settings employed in this study have been previously described ^114^.

### cDNA synthesis and library preparation

Gel Beads in Emulsion (GEM) generation, cDNA synthesis and library construction were carried out as indicated by the 10× Genomics Chromium Next GEM Single Cell 3ʹ Reagent Kits v3.1 user guide. For each sample, 10× microfluidic chips were loaded with 10 thousand nuclei, and 13 PCR cycles were used for cDNA amplification. Subsequent cDNA library sequencing was carried out on the NovaSeq 6000 System at the ICBR, with S1 flow cell 2x100 sequencing kit but with cycling of 28 read 1, 10 index 1, 10 index 2, and 90 read 2. For the Dusk Salt sample, a second technical replicate was obtained by re-sequencing of the library a second time to obtain additional reads and equilibrate the total number of nuclei identified in each sample. Both sequencing files from the same Dusk Salt library were concatenated prior to downstream analyses.

### Data processing and clustering

Data processing steps were carried out on HiperGator. The cDNA sequences were processed and the counts matrix was generated using Cell Ranger (v7.0, 10× Genomics) with default parameters. For each sample, an expected number of 10 thousand cells were considered using the option “-- expect-cells”. The raw data from Cell Ranger was used for subsequent quality control and filtering steps. First, we took measures to eliminate potential contamination from chloroplast cDNA. To achieve this, we removed genes with high similarity (BLASTN, e-value < 1x10^-8^) to the ones encoded by the *M. crystallinum* chloroplast genome (NCBI NC_029049) from the raw counts matrix (**Supplemental Table 11**). To identify empty droplets and low-quality cells, we used the emptyDrops function from the DropletUtils package (version 1.20.0). We ensured the appropriateness of the model for removing empty droplets by testing it on each sample, as per the recommendations in the package documentation, using the “test.ambient=TRUE” option. Under the null hypothesis, p-values for empty cells (cells with total counts below the set threshold) should distribute uniformly. Therefore, we applied various UMI count thresholds to each sample until we achieved a uniform distribution of p-values for cells with low total counts. This approach allowed us to determine a unique minimum UMI threshold for each sample, effectively removing low-quality nuclei while preserving biologically relevant nuclei. Details of the thresholds applied, and the nuclei retained can be found in **Supplemental Table 12**. Any gene that was not detected in at least three cells was excluded from the Seurat objects. Data pre-processing and analysis steps, such as normalization, and batch-effect removal, were conducted using Seurat V4.3.0. Integration and clustering of the four snRNA-seq datasets was carried out at a resolution of 0.55. Uniform Manifold Approximation and Projection (UMAP) was used to reduce the complexity and visualize the data in two dimensions. Thirty components were consistently selected for dimensionality reduction across all samples. To enhance the accessibility and exploration of our dataset, we developed an interactive, user-friendly online Shiny application. This application was created using the ShinyCell R package (version 2.1.0). It offers a convenient platform for users to engage with and analyze the data. The app is publicly available and can be accessed at: https://noeperron-kirstlab.shinyapps.io/shinyapp/

### Gene orthology and annotation of *M. crystallinum* cell-type-specific markers

Gene orthology throughout this study was determined by aligning protein sequences to the Swiss-Prot database using BLASTP with a stringent e-value cutoff of 1e-6. Orthologs were identified based on the reciprocal best-hit method, with the top hit considered the ortholog of the ice plant gene. In cases where multiple genes exhibited similarly high levels of similarity, both were designated as orthologs. Cell-type identification was conducted by analyzing the expression profiles of orthologs to known cell-type specific markers from the literature in the ice plant genome (**Supplemental Table 3**). Furthermore, cell-type assignments were refined by examining the known functions of orthologs expressed within distinct clusters. This was done using Seurat’s FindMarkers() function, which performs differential expression testing between clusters using the Wilcoxon rank sum test. Genes were classified as differentially expressed if they had a log2 fold-change greater than 1 and an FDR below 0.05 (**Supplemental Table 4**).

### hdWGCNA analysis

Co-expression networks were constructed on the integrated snRNA-seq dataset using the hdWGCNA R package ^84^. Only genes expressed in at least 5% of cells were retained for this analysis. Metacells (or groups of cells with similar transcriptomic signatures) were constructed by grouping cells based on Seurat clusters using PCA for K-nearest neighbors (KNN) analysis with k=25 and a maximum of 10 shared cells between metacells. The metacell expression matrix was normalized, and the expression data for each cluster was prepared. Various soft-power thresholds were tested to determine the optimal value for network construction, and a signed co-expression network was constructed using a soft-power threshold of 7 as determined by the algorithm. Ten modules were identified, and eigengene-based connectivity (kME) was calculated for each module to rank genes by order of connectivity to their respective module.

### Tissue collection for bulk RNA-sequencing

*M. crystallinum* plants were grown under the conditions described in the “Plant Material Growth Conditions for snRNA-seq” section. Four-week-old plants were watered daily with 50 mL of 0.5X Hoagland’s solution. To induce CAM, a subset of plants was watered with 50 mL of 0.5X Hoagland’s solution containing 0.5M NaCl. After 21 days of salt treatment (seven-week-old plants), the third pair of leaves was collected from both the well-watered and salt-treated groups every four hours from 8:00 AM (ZT0) to 4:00 AM (ZT20) the following day, totaling six time points. Three biological replicates were harvested per condition, resulting in 36 samples.

### RNA extraction and sequencing

Total RNA was extracted from the 36 samples using a modified version of a previously described protocol ^109^. Samples ground to a fine powder were transferred into 500 µL of warm CTAB extraction buffer with added 2-mercaptoethanol. An equal volume of chloroform was added, and phases were separated by centrifugation at 10,000 rpm at room temperature. An equal volume of 7.5M LiCl was added, and RNA was precipitated overnight at 4°C and harvested the next day by centrifugation at 10,000 rpm at room temperature. After two ethanol washes, RNA was resuspended in an elution buffer. RNA was subsequently cleaned using an RNA Clean & Concentrator kit (R1016, Zymo Research, USA) following the manufacturer’s protocol. cDNA libraries were constructed using the NEBNext® Ultra™ Directional RNA Library Prep Kit for Illumina® (NEB #E7760) according to the manufacturer’s instructions. Paired-end sequencing was carried out on a single NovaSeq 6000 System lane at the ICBR, with a 100-bp read length and a sequencing depth of over 1 billion reads.

### Data processing and identification of DE genes

RNA-seq reads were processed starting with the trimming of adapters and low-quality bases using Trimmomatic (v.0.39). After trimming, the quality of the reads was assessed using FastQC (v0.11.9). The trimmed reads were aligned to the *M. crystallinum* reference genome presented here using HISAT2 (v.2.2.1). Gene expression was quantified using HTSeq-count (v.2.0.3), which provided the counts of reads mapping to each gene. The gene count matrix generated by HISAT2 and HTSeq-count was loaded into R for differential expression analysis using the edgeR package (v. 3.40.2). To ensure robust statistical analysis, genes with low counts were filtered out, retaining only those with a 75th percentile count greater than or equal to 5 in at least 25% of the samples. Normalization was performed using the TMM (trimmed mean of M-values) method to account for compositional biases between libraries. Differential expression testing was conducted using an exact test to compare counts between salt-treated and well-watered samples. Top differentially expressed genes were identified and filtered by false discovery rate (FDR, **Supplemental Table 10**).

### Time-series gene co-expression network inference

Time-series gene co-expression network inference was performed on the bulk RNA-seq data using the maSigPro R package (v.1.70.0) ^86^. The raw count matrix was normalized using the “calcNormFactors” function from the edgeR package (v. 3.40.2), followed by a transformation using the “voom” function from the limma package (v. 3.54.0) to stabilize the variance-mean relationship in the data. We constructed a design matrix for maSigPro with a polynomial degree of two, specifying the columns for time, replicates, and experimental groups. Significant genes were identified using the “p.vector” and “T.fit” functions, and the “get.siggenes” function was employed to summarize these significant genes. To determine the optimal number of clusters for the significant gene profiles, we calculated the total within-cluster sum of squares (WSS) for 1 to 15 clusters using k-means clustering and plotted an elbow plot. The optimal number of clusters was chosen based on the point where the reduction in WSS started to slow down significantly, which in this case was four clusters.

### *In vivo* transactivation assays

To assess the ability of HY5 and HYH transcription factors to activate PPCK1 expression, transactivation assays were performed in *Nicotiana benthamiana* leaves. Promoter sequences corresponding to *PPCK1.1* and *PPCK1.2* from *M. crystallinum*, as well as the *AtPPCK1* promoter from *Arabidopsis thaliana*, were amplified from genomic DNA and cloned into a LUCIFERASE (LUC) reporter construct (LUC::Promoter::tNOS) using the GAANTRY cloning system^115^. Effector constructs (35S::TranscriptionFactor::tNOS) encoding McHY5, McHYH, AtHY5, or AtHYH, and reporter constructs (35S::GUS::tNOS) were similarly cloned. *Agrobacterium*-mediated transient expression was performed by infiltrating *N. benthamiana* plants (4-6 weeks old) as previously described^116^ with strains carrying the reporter, target, and effector constructs. Control leaves were infiltrated with *Agrobacterium* carrying only the reporter and target constructs. Leaves were harvested three days post-infiltration for LUC and β-glucuronidase (GUS) activity measurements. Five biological replicates were performed for each effector/reporter combination. Statistical significance between effectors and the control was determined using a Wilcoxon test, with significance thresholds of p < 0.05 (*) and p < 0.01 (**) (**Supplemental Table 13**).

## ACKNOWLEDGMENTS

The authors extend their heartfelt gratitude to Dr. Klaus Winter from the Smithsonian Tropical Research Institute for his invaluable contributions to this manuscript. His expert insights on CAM significantly enhanced the quality of our work. The authors also thank Mariza Miranda from the Flow Cytometry and Confocal Microscopy core at the Interdisciplinary Center for Biotechnology Research (University of Florida). Finally, the authors thank Dr. Sixue Chen from the University of Mississippi Department of Biology for providing *M. crystallinum* seeds.

## AUTHOR CONTRIBUTIONS

Conceptualization, NP and MK; Software, NP; Formal Analysis, NP; Investigation, NP; Data Curation, NP; Visualization, NP; Writing – Original Draft, NP; Writing – Review & Editing, NP, WJP and MK; Methodology, NP, CD, WJP, and MK; Resources, MK and BB; Supervision, CD and MK; Project Administration, NP, CD, and MK.

